# Gene-Culture Coevolution of Prosocial Rituals

**DOI:** 10.1101/060632

**Authors:** Karl Frost

## Abstract

It has been argued that costly socially learned rituals have the potential to generate prosocial emotional responses arising from genetically based behavioral dispositions, hijacking these behavioral dispositions to solve otherwise intractable cooperation problems. An example is the innovation of synchronous movement rituals which seem to engage the same altruistic response as instinctual mimicry. Arguments against this hypothesis are based in the idea that cheap markers of altruistic intent (“Green Beards“)are vulnerable to manipulation from non-cooperating free-riders that are similarly marked. This paper formally models the gene-culture coevolutionary dynamics of a system in which a costly socially learned ritual practice coevolves with the genes underpinning the prosocial response. This takes a ‘genetic mismatch’ hypothesis (where fast evolution of socially learned behaviors create a temporary mismatch with the coevolving genes, which take longer to evolve to equilibrium) and models it dynamically. The genetic and cultural fitness equations are first analyzed for equilibrium and then simulations are used to predicting trajectories for both the socially learned behavior and the hijacked genetic trait over time. Relying on fast culture and slow genes, it demonstrates how high levels of ritual efficacy may be established, at least temporarily, through gene culture interaction. It also shows how longer term feedbacks on the genes from culture may lead to decreases in the prosocial gene in the population and reduced overall population mean fitness, despite high levels of altruism in the population.

## Introduction

Many rituals are practiced with the intention of bonding together people into cooperative groups, willing to sacrifice for each other and the benefit of the group (Durkheim, 1912; Rappaport, 1999). These rituals are often costly in terms of time investment and energy. Many researchers currently focus on rituals involving intentional, synchronized or coordinated movement and propose that such rituals regularly evoke altruism amongst co-participants (Kirschner & Tomasello, 2010; McNeill, 1995; Reddish, Fischer, & Bulbulia, 2013). This response to these socially learned practices is suggested to be based in instinctual mimicry actions and the altruistic response that such generates (Weingarten & Chisholm, 2009). In more general terms, this proposes that a socially learned ritual practice can hijack a pre-existing sensory bias to create altruistic behavior in new contexts. This then sets the stage for gene-culture coevolutionary dynamics. In this light, it also provokes criticism as an hypothesis, given how similar it is to a Green Beard proposal. A Green beard is where a marker is associated with altruistic response and thus serves as a means for altruists to preferentially associate with each other to overcome problems of non-reciprocating free-riders taking advantage of their altruism (Dawkins, 1976; Hamilton, 1964). This proposal is recognized as problematic, as the system is evolutionarily vulnerable to invasion from individuals who have the marker but not the altruistic response. The evolutionary stability of altruism in green beard models relies on some mechanism maintaining sufficient covariance between the marker and the altruistic response. In this paper, I formally model the proposal of costly socially learned rituals hijacking pre-existing behavioral responses to induce altruism using gene-culture coevolution models. I first solve analytically for equilibrium trait frequencies and then simulate gene-culture coevolution to predict trajectories of trait frequencies over time and, by corollary, the time scale of change in frequencies. I show that if the ritual is costly and/or that there is a pre-existing benefit to the hijacked allele then more complex coevolutionary dynamics can happen than in the case of a simple Green Beard model, leading to long term altruism in the population. If the preexisting benefit of the hijacked allele is sufficiently strong, the ritual behavior moves to fixation as does ritual facilitated altruism. In the intermediate case where the benefit of the hijacked allele is less than the cost of cooperation in Prisoner’s Dilemma (PD) type situations and where the cost of the ritual is less than net benefit to be found through cooperation in a PD in a population of altruistic cooperators, then the population moves through damped oscillations in frequency space of traits around an unstable equilibrium point set by these parameters. This damped oscillation lasts a long time compared to the case where the population moves to fixation. In the short term, as on the scale of human history, the trajectory of such oscillations may be characterized by relative fixation in ritual facilitated altruism before the population moves further through the oscillations to eventual decline in altruism. This dynamic is set up by the difference in time scales of genetic vs cultural change (Perreault, 2012). In another paper, I look at the genetic rather than cultural evolution of ritual behavior (Frost, 2016a), and the speed of change is central to the differences in predictions of the two models.

## Ritual proposed as reliable signal of altruistic intent

Several proposed mechanisms can promote and stabilize altruism in groups of unrelated individuals. Repeated play and reciprocal altruism can do this (Trivers, 1971)(Axelrod & Hamilton, 1981). Cheap punishment reinforced by conformist social learning and norm psychology can also bolster cooperation and altruism (Henrich & Boyd, 2001). Still, there is the problem of what we do when no one is watching. Neither punishment nor reputation support altruism when cooperators and defectors can’t be identified. One can imagine situations in which punishment or reciprocity can affect ‘invisible behaviors’ when those behaviors have subtle cues or when there is occasional visibility and strong enough punishment to compensate for the infrequency of punishment. Yet, what about the cases where one is certain that one will not be caught defecting? Most would expect that people are less likely to be altruistic in these instances, yet it is also clear empirically that altruism does not go away and remains an important element of human social relations even when not enforceable (Batson, 2011). If there are behaviors that could reliably increase discreet altruism, this would be of enormous benefit to groups. If there was some means to establish positive assortment of such discreet altruists, it could be maintained through group selection (Sober & Wilson, 1998). A reliable marker associated with altruistic intent could allow for positive assortment amongst altruists. This is the classic ‘green beard’ solution: if altruists and only altruists had green beards, assortment based on beard color would result in groups of only altruists which could then flourish through group selection (Hamilton, 1964) (Dawkins, 1976). While such markers have been shown to exist, it has been argued that they are likely to be uncommon, as genetic recombination would tend to lead to the breaking of linkages between the genes for the marker and the genes for altruism (Grafen, 1998).

Cultural evolution models demonstrate that arbitrary socially learned ethnic markers can facilitate the spread of coordination through cultural group selection (McElreath, Boyd, & Richerson, 2003). However, the evolution of *altruism* is not predicted by this model, and, being based on social learning, it is problematic using this as an explanation of discreet behavior that is, by definition, not witnessed and therefore difficult to socially learn(Frost, 2016b). The mechanism of costly rituals is novel, using the hijacking of a genetic behavioral trait by a socially learned behavior (the ‘ritual’) to generate altruism that does not rely on the visibility of cooperation/defection.

I present a simple mathematical model showing that such hijacking, in the form of ritual behavior, can arise and persist. This argument is not novel, as the idea of rituals engaging genetically evolved behavioral dispositions is part of the basis of a number of theories of ritual (Frost, 2013; McNeill, 1995; Swann, Jetten, Gómez, Whitehouse, & Bastian, 2012; Tomasello, Melis, Tennie, Wyman, & Herrmann, 2012; Weingarten & Chisholm, 2009). What is unique in this paper is the formal modeling of coevolution of the traits and the novel contributions of the differences in genetic and cultural change. Further the verbal arguments of gene-culture mismatch generally assume that genes do not evolve in timescales of human cultural history. The models presented here explicitly models these dynamics, generating novel predictions for the frequency of rituals and the genes they hijack over time, and generating a more complex picture of such hijacking.

Many have argued that behavioral mechanisms inherited via genetic selection trump and canalize socially learned behaviors, keeping socially learned behavior on a tight leash (Lumsden & Wilson, 1981). However, such genetically evolved leashing, should it exist, would be evolutionarily shaped by historical, not new circumstances. Inasmuch as we can use assumptions of adaptation, we can not assume adaptation to circumstances that have not yet occurred. Part of the purpose of this paper is to formally model the gene-culture coevolutionary dynamics to reveal what might be thought of as a ‘leashing effect,’ yet are more complex than those arising from verbal arguments informed by genetic equilibrium assumptions. Where the culturally evolved behavior (the ‘ritual’) is visible, and the triggered genetically determined behavior is prosocial in the novel context, the socially learned ritual behavior becomes a reliable visible signal of intent to reciprocate cooperation. The ritual takes advantage of the genetically evolved behavior, and the slow evolutionary response of genes creates a long term changing genetic mismatch between the trait frequencies and genetic fitness optimization.

## Synchrony

While the modeling I do here of reciprocal ritual is more general, it is based on the empirical example of intentional synchronized movement. Ethnographic and historical evidence supports the notion that group unison movement facilitates a sense of group and a willingness to sacrifice for the benefit of fellow group members (McNeill, 1995). There is also growing experimental evidence demonstrating the in-group prosocial benefit and group forming capacity of synchronized movement (Kirschner & Tomasello, 2010; Reddish et al., 2013) and mimicry (Chartrand & Baaren, 2009). The historical ubiquity of these rituals and the ability to consistently trigger such prosocial responses suggests a genetic basis, rather than the effect of socially learned and culturally flexible norms. Weingarten and Chisholm (2009) have suggested that the evolutionary roots of this instinct are in the increased importance of infant-caregiver bonds in an environment of extended childhood development, while Tomasello et al (2012) suggest that the roots of this prosocial response are in obligate collaborative foraging. The evidence for such a widespread human prosocial synchrony instinct is, in any case, compelling.

Once rapid, cumulative social learning (culture) evolves, it sets the stage for hijacking of instinctual prosocial responses to mimicry, activating these responses ‘artificially’ in the novel context of socially learned ritual performance. Should anew habit of performing unison movement be innovated, the behavioral tendency to be altruistic would be activated within these groups of unison movers. This ritual, spread through social learning, would then improve cooperation in these groups and help them overcome anonymous prisoner’s dilemma style cooperation problems.

Even in a situation where the stakes are large in the PD game, and the free rider problem is severe, it may be possible that the relative speed of social learning in comparison to genetic change allows, at least for the ‘short term’ of recorded human history, for cultural hijackers to sweep to fixation in a population. Gene frequency would very slowly respond with an increase in free-riders psychologically unresponsive to the ritual they perform. The change in the genetic composition of the overall population might be imperceptible in the short term of centuries, but perhaps be significant in our longer evolutionary history.

## The model - Costly ritual as mutual manipulation

The model presented here is a simple one. Actual evolutionary dynamics are undoubtedly more complex. Single genes coding for complex social behaviors seem improbable, as do socially learned behaviors without a wide range of variants. Simple models, however are useful to qualitatively demonstrate evolutionary dynamics (Boyd & Richerson, 1985; Levins, 1966; Winterhalder, 2002), which is the purpose of this paper. In this section, I formalize the verbal model mathematically and explore coevolutionary equilibrium analytically. In the next section, I use simulations to explore the trajectory of the gene frequencies in the population over time.

I model the gene-culture coevolutionary dynamics of a socially learned reciprocal ritual behavior that hijacks an existing sensory bias to induce altruistic behavior. It is identical to the purely genetic coevolution model I explore in a separate paper, except that the ritual, rather than being genetically determined, is acquired through success-biased social learning (Frost, 2016a). The ritual is reciprocal in the sense that it involves participation of both individuals and can not be performed unilaterally. A person can not perform a synchronous dance unless they have a partner with whom to synchronize. Assume a system with a single gene and a cultural trait. Both the gene and the cultural trait have two variants.

The gene codes for the contextual altruistic behavior; allele *A* has this contextual altruistic response to ritual and allele *a* does not. Allele *A* also has a simple additive fitness benefit, g, over allele *a.* In our example, A causes improved mother child bonding, and g is this benefit. This can be thought of formally like a pleiotropy, though it is not actually two separate response, but the same response activated in two different circumstances by similar stimuli. With all else being equal, *A* would move to fixation in the population, and *a* would only exist in the population as a vestige or rare mutation.

**Table 1:** Encounter outcomes by allele combinations: PD play (Ego/Other) and net fitness benefit (Ego). In PD play, c refers to altruistic cooperation and d refers to defecting/free-riding

|  |  | Other |  |  |  |  |  |  |  |
| --- | --- | --- | --- | --- | --- | --- | --- | --- | --- |
|  |  | AE |  | Ae |  | aE |  | Ae |  |
|  |  | PD Play<br>(Ego/Other) | Fitness<br>benefit<br>(Ego) | PD Play<br>(Ego/Other) | Fitness<br>benefit<br>(Ego) | PD Play<br>(Ego/Other) | Fitness<br>benefit<br>(Ego) | PD Play<br>(Ego/Other) | Fitness<br>benefit<br>(Ego) |
| Ego | AE | c/c | $g+(b-c)-f$ | d/d | $g$ | c/d | $g-c-f$ | d/d | $G$ |
| | Ae | d/d | $g$ | d/d | $g$ | d/d | $g$ | d/d | $G$ |
| | aE | d/c | $b-f$ | d/d | $0$ | d/d | $-f$ | d/d | $0$ |
| | ae | d/d | $0$ | d/d | $0$ | d/d | $0$ | d/d | $0$ |

There are two cultural variants with regards ritual performance; variant *E* performs the ritual with willing others (other Es), while variant *e* never performs the ritual. When two individuals meet they are faced with two sequential choices: firstly, whether or not to perform a ritual, at cost f, and subsequently how to behave in a Prisoner’s Dilemma (PD) type situation. The PD has a cost, c, to cooperate. If one cooperates, one’s partner gets benefit, b. Consider our synchronous movement example. f in this case would be the cost of time and energy to engage in the dance. c would be the choice to work extra hard in some shared project, to take greater risks that would benefit the other, or perhaps to be trustworthy in economic activities. b would be the benefit to others from these altruistic acts. f<(b-c). Cooperation is the altruistic choice. Defection is the selfish choice to not put in any investment that is not immediately of net benefit to the self. I assume that cooperation/defection are not visible, so there is no opportunity for reputation or reciprocity to affect strategies. The evolutionary equilibrium play for such simple PD games in the absence of ritual sensory hijacking is to defect.

**Table 2:**
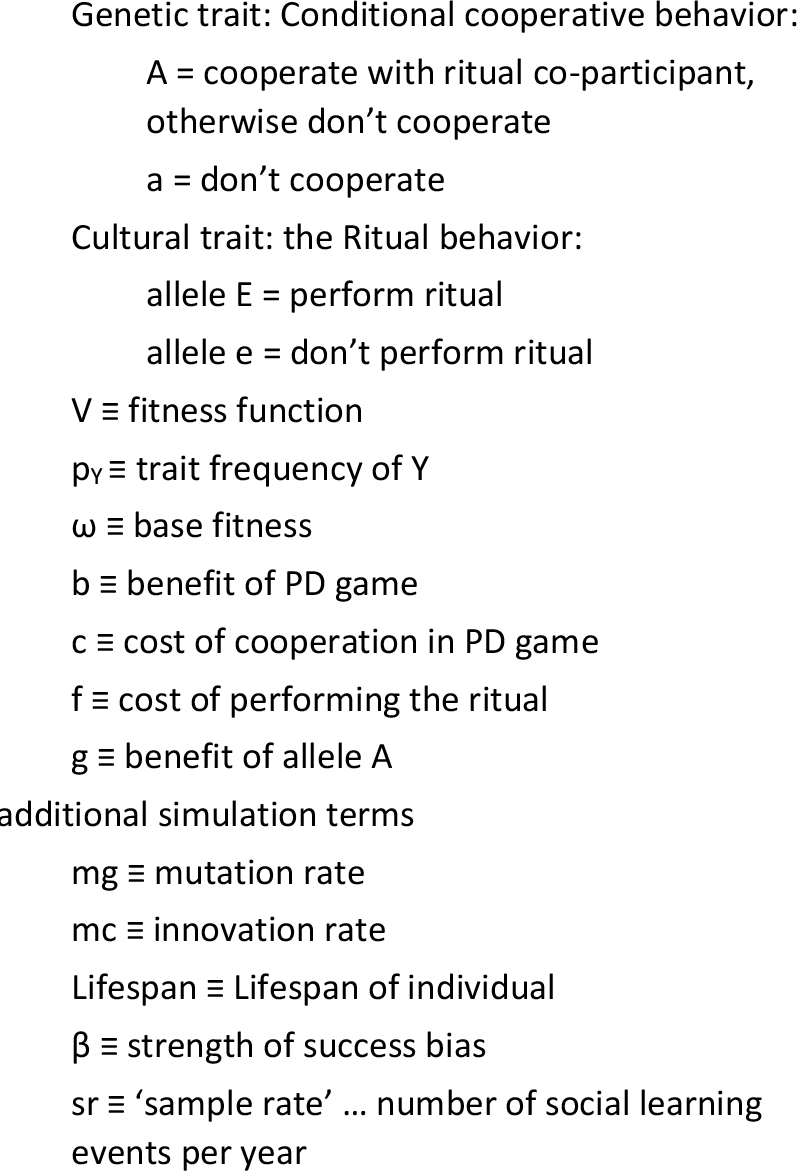
Equation terms

*Ae* does not perform ritual and defects. *AE* is willing to perform the ritual dance, and cooperates in PD with any ritual co-participant, if they find one. *aE* is willing to perform ritual but defects with all, even with ritual co-participants. *ae* does not perform ritual and likewise defects.

Ae and AE both get the benefit, g, of improved mother/child bonding. If an AE or aE meets another E, they perform the dance at cost f. An AE performing such a dance will cooperate with their partner, whether AE or aE. aE engages in ritual dance (at cost f) but defects in PD situations, motivated by their own personal fitness. aE will defect on their dance partner, but, of course, they also lack the benefit, g, of superior mother/child bonding. The gene for A/a is invisible, so ritual participants do not ‘know’ a priori if they will receive cooperation or defection. In the absence of E, there is no ritual performance and no cooperation. In the absence of A, there is no cooperation, even if there is ritual performance.

Table 1 gives the plays in the PD Game and the fitness of allele combinations based on encounters. Table 2 summarizes the fitness equation terms as well as parameters for subsequent simulations.

Population mean fitness values of the different allele combinations, with ω as base fitness are

- V (ae) = ω
- V (Ae) = ω + g

In the case where there are no E variants in the population, the population tends to move to p_A_ =1, where p_Y_ is the fraction of the population with allele Y, and p_XY_ is the fraction of the population with allele combination XY. With E variants, we have

- V (AE) = ω + g + p_AE_b – p_E_c – p_E_f
- V (aE) = ω + p_AE_b – p_E_f

Assume haploid genetics, reproduction in proportion to the fitness value. Social learning is success biased: with additive chance of copying a behavior from another proportional to the difference in genetic fitness between self and other. Assume thorough recombination between cultural and genetic variants.

## Results

Doing a little algebra, I find the equilibrium values of p_A_ and p_E_ given in Table 3. The difference in population mean fitness between equilibrium and a population of all Ae is also given. This is taken as the starting population in simulations in the next section. These equilibrium results are identical to those in the case of genetically based ritual. behavior.

**Table 3:**
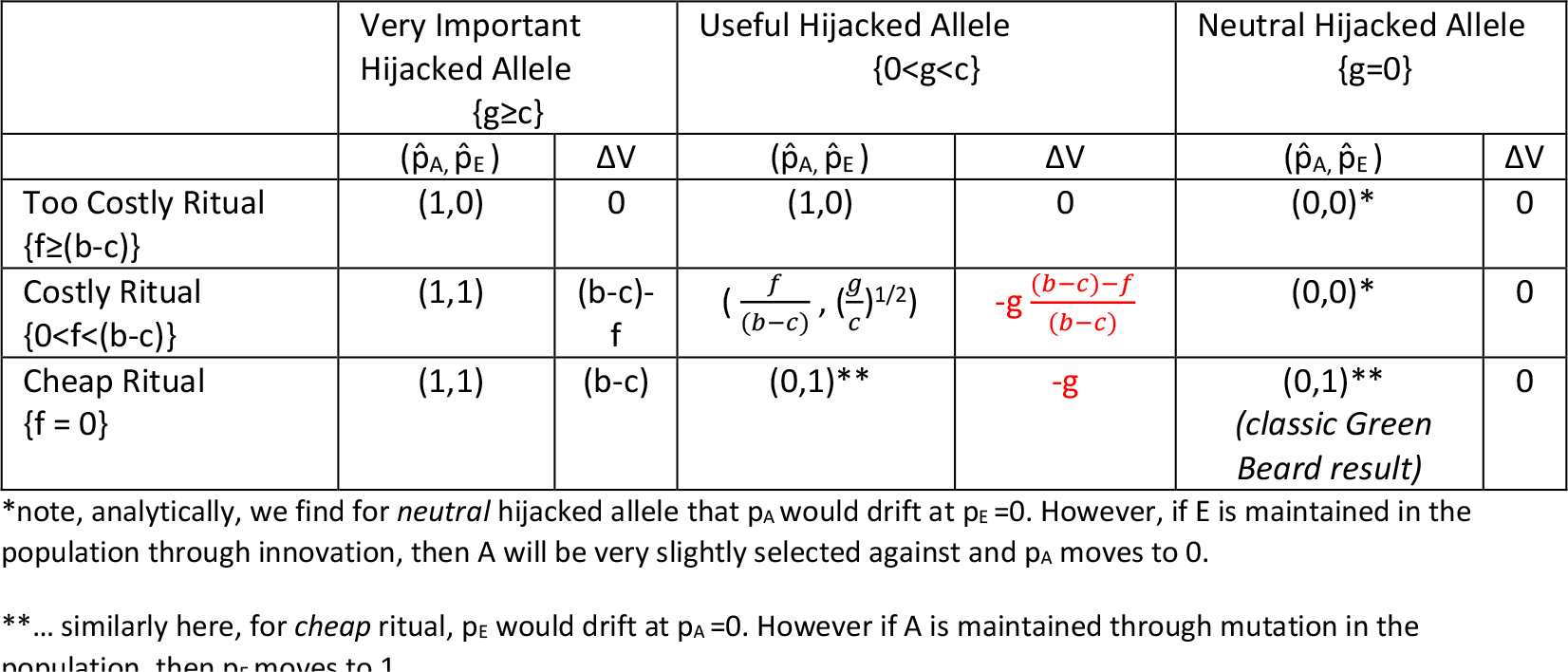
Equilibrium for different parameter combinations: gene frequencies 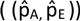and change in population mean fitness from all Ae to equilibrium (ΔV)

There are 3 relevant ranges of parameters for the ritual and hijacked allele. For the ritual, we can call these *cheap* (f=0), *costly* (0<f<(b-c)), and *very costly* (f≥(b-c)). For the hijacked allele, we can call these *neutral* (g=0), *useful* (0<g<c), and *very useful* (g≥c). There is some parameter combination that would keep any one of the allele combinations at fixation.

The case of *cheap* ritual (f=0) and *neutral* hijacked allele (g=0) matches the conventional green beard model and replicates the results of such. In this case, there is no extra mother/child bonding benefit and the ritual triggering the altruistic response is cheaply done. Altruism does not survive in the population at equilibrium, as is the prediction in the simple green beard model. We would expect ritual facilitated altruism to rise sharply at first, but then free riders do very well and eliminate the altruistic allele. This is demonstrated in simulations in the next section. Interestingly, the ritual is ubiquitous and thus the potentially altruistic gene is kept out of the population without some countervailing influence. The population mean fitness in the end is unchanged as it rises and then goes back down at equilibrium after the introduction of the ritual gene. Interestingly, the same result is achieved even if the hijacked allele was *useful* (but not when *very useful*): A is eliminated by ubiquitous cheap rituals. Here, the population mean fitness is reduced by g, the benefit of the hijacked allele. Of course, when the ritual is *very costly*, it fails to evolve in all cases.

When the hijacked allele is *very useful* it is maintained at fixation in the population, barring rare mutants. In this case, if the ritual is *costly* or *cheap* it also moves to fixation in the population as does ritual-facilitated altruism. This is reflected in the increase in population mean fitness in the amount of the net benefit of 100% cooperation in the prisoner’s dilemma, minus the cost of the ritual: ΔV = (b-c) − f.

The case of intermediate parameter values-*costly* rituals and *useful* hijacked alleles (c>g>0 and (b-c)>f>0)-results in a balance of all 4 allele combinations at equilibrium.

- For V(A) = V(a), 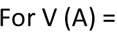
- For V(E) = V(e), 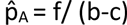

Figures 1A and 1B show these lines of reproductive value equivalence and the direction and magnitude of selection on the traits for different points in trait frequency space for such a case. The equilibrium frequency of the hijacked allele is set by the ratio of the cost of the ritual and the potential net benefit for two altruists in the PD game. Counterintuitively, this equilibrium frequency of the hijacked allele *does not* depend on how beneficial the hijacked allele is on its own (g), so long as 0<g<c. As the cost of ritual performance (f) increases, the equilibrium frequency of altruistic behavior increases. However, the evolution of the ritual behavior results in a *reduction* of population mean fitness at equilibrium, unless g>c. The *costly/useful* equilibrium is unstable, however. I show in simulations that the population cycles through damped oscillations of trait frequencies, influenced very strongly by the rate of innovation, with a generally declining population mean fitness, from starting point at fixation in Ae. The selection pressures illustrated in Figure 1 give an intuition for why this cycling would be the case. The decrease in population mean fitness of the population happens despite the evolution of altruism in the population, due to the combined effects of freeriding, the reduction of the frequency of the beneficial hijacked allele, and the costly investment in ritual.

**Figure 1a.**
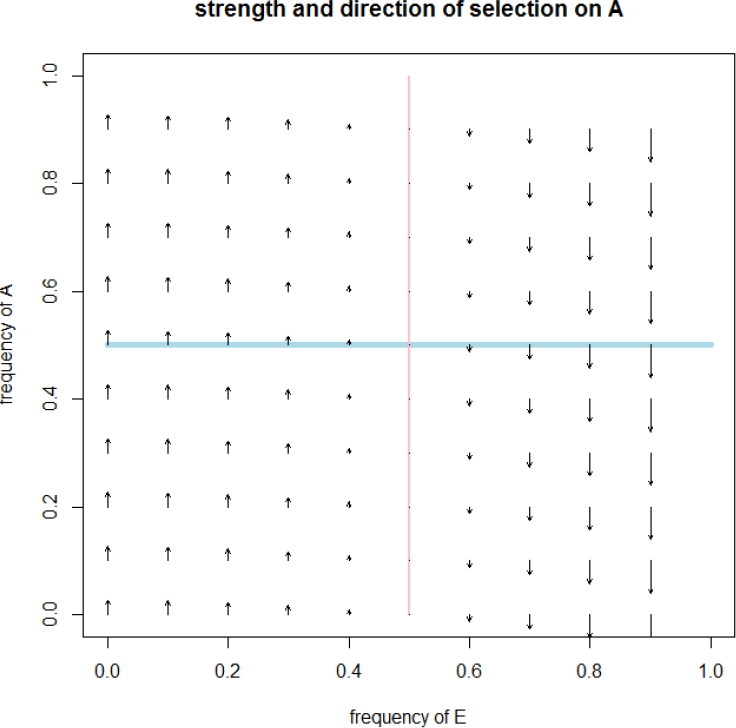
strength of selection on A for intermediate parameter values {*g* = .125, *b* = 1, *c* = .5, *f* = .25}

**Figure 1b.**
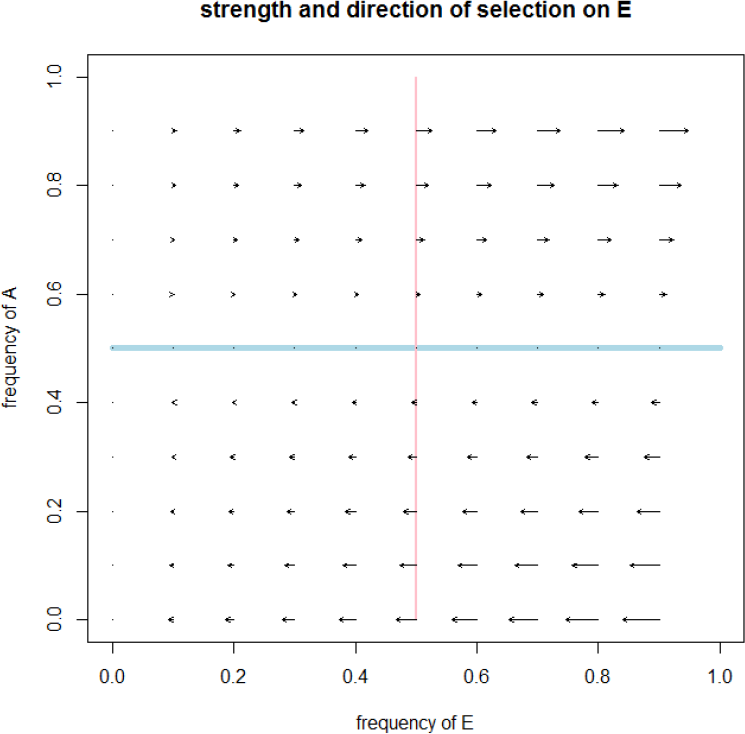
strength of selection on E

*Note, these analytic results and the simulations below are based on an assumption of thorough recombination between genes and cultural variants in a well-mixed population. Covariance is assumed to be non-existent. While covariance is often small enough to have negligible effect in evolutionary models, especially in a socially learning population, sometimes even very small amounts of covariance can have dramatic effects. In an appendix, I share analysis where recombination is explicitly modeled via social learning and where covariance is allowed to remain. I show that the dynamics are almost indistinguishable from the case of complete recombination. See Appendix on recombination.*

These analytic results for equilibrium are the same as if the ritual behavior were genetically determined rather than acquired through success-biased social learning. Analysis of the genetic model is shared in a separate paper (Frost, 2016a). Differences emerge in simulations based on the relative speed of cultural vs genetic change.

## Simulations

At the start of simulations, the population is entirely Ae, having the mother/child binding gene, but lacking ritual performance. As the simulation progresses, I assume a well-mixed population. Each year, individuals of a given genotype reproduce proportional to the ratio of their fitness to base fitness, divided by Lifespan. Newborns are subject to mutational change from one allele to another (*m*= 2.5 × 10^−5^ changes/generation).

The population engages in one social learning event per year. Social learning of E or e is success biased, with individuals more likely to learn from individuals with higher demonstrated genetic fitness. In each social learning event, an individual randomly meets another and engages in success biased social learning, where their chance of changing their behavior to that of the encountered other is given by

> Chance of acquiring cultural variant of other via social learning = 0.5 + β ΔV

where β is a scalar of the strength of social learning bias and ΔV is the difference between the fitness of the observed other and the fitness of the self. This form of success bias is commonly used in economic and cultural evolution models (McElreath & Boyd, 2007). Again, there are sr = 1 social learning events per year. As sr or β increase, the speed of cultural change would increase. In each social learning event, there is also a chance of innovation. Paralleling mutation, innovation is a spontaneous change of cultural variant, the chance of which is given by the innovation rate, set at 1/10000 per social learning event.

Lifespan (50 years), base fitness (*w* = 10), PD benefit (*b* = 1) and cost (*c* = .5) are held constant amongst simulations. Simulations were run for 150,000 years. Benefit of hijacked allele (g) and cost of ritual (*f*) are varied to represent the hijacked allele being *very important* (g>c … g =1), *useful* (c>g>0 … g =.125), or *neutral* (g=0), and to represent the ritual being *cheap* (f=0), *costly* ((b-c)>f>o … f = .25), or *too costly* (f>(b-c) … f = 1). The simulations begin with initial conditions, p_A_ = 1, p_E_ = 0. This represents the condition where A arose in our deeper past and E arose in the context of more recently evolving sophisticated social learning skills. Genetic and cultural variants are introduced through mutation and innovation. See appendix for R code.

## Simulation Results

Simulation results are illustrated in Figure 2. Figure 2A shows the trajectory through the space of trait frequencies, Figure 2B the genetic and cultural variant frequencies over time, and Figure 2C the population mean fitness over time. The simulations support the analytic results for equilibrium as well as give a qualitative sense of the dynamics over time. The pathway over time is important for comparison to timescales of human history, especially relevant for intermediate values of f and g.

Some observations from the simulations.

- For f<(b-c) (*costly* or *cheap* rituals, but not *too costly),* the frequency of *E* increases rapidly at first, moving to near fixation within a couple of hundred years. It then stays near relative fixation until the more slowly moving gene frequency catches up through coevolutionary processes, in the case of *useful* or *neutral* hijacked alleles.
- Where allele A is *very useful (g>c), A* stays at relative fixation as does E. All individuals eventually perform moderately costly or cheap rituals and cooperate with ritual co-participants.
- For g<c, the response of genes to the advent of widespread ritual use happens on the order of 20k years for *g<c.* These alleles decreasing in frequency, due to the advantages of free-riding.
- As expected, prohibitively costly rituals (f>(b-c)) do not evolve and are only maintained in the population as a result of random experimental innovation. The simulations used a rate of random innovation at 1/10000 seemingly small, but in the simulations sufficient to maintain the ritual in the population at a rate of approximately 5% even when otherwise too costly. For more borderline cases, this could have siginificant impacts on gene culture coevolution.
- For f=0 and g<c, the population moves toward fixation in E and a, which again mirrors the findings for green beards.
- The case of neutral hijacked allele and costly ritual does not in the time frame of the simulations move significantly toward the equilibrium found analytically. However, in closer inspection, it is found that the populations is moving toward p_ae_ = 1. It is just happening very slowly, owing to the quite miniscule selection pressures involved. This was verified in a series of side simulations.
- For the intermediate case of *useful* hijacked allele and *costly* ritual, the trait frequencies can be described as having 4 phases. In the first, there is rapid swing to fixation of ritual practice. This takes a few hundred years. The next phase, lasting about 20,000 years, has the hijacked allele A, near fixation but the alternate allele, a, slowly rising in frequency. The next phase, also lasting of the order of 20,000 years, has the genes and culturalvariants moving through damped oscillations toward equilibrium. As culture responds much more quickly to selection pressures than genes, the frequency of E varies more widely around equilibrium than does the frequency of A. The final phase, which goes on indefinitely, has the population very close to equilibrium with quite small damped oscillations around it. The damping of the cycles is very strongly influenced by innovation rate, just as the cycling in the purely genetic case is influenced by mutation. Higher innovation rates would dampen the cycles more, causing the population to spiral farther in toward a limit cycle closer to the (unstable) equilibrium. This can be demonstrated by running simulations with varied innovation rates.

**Figure 2:**
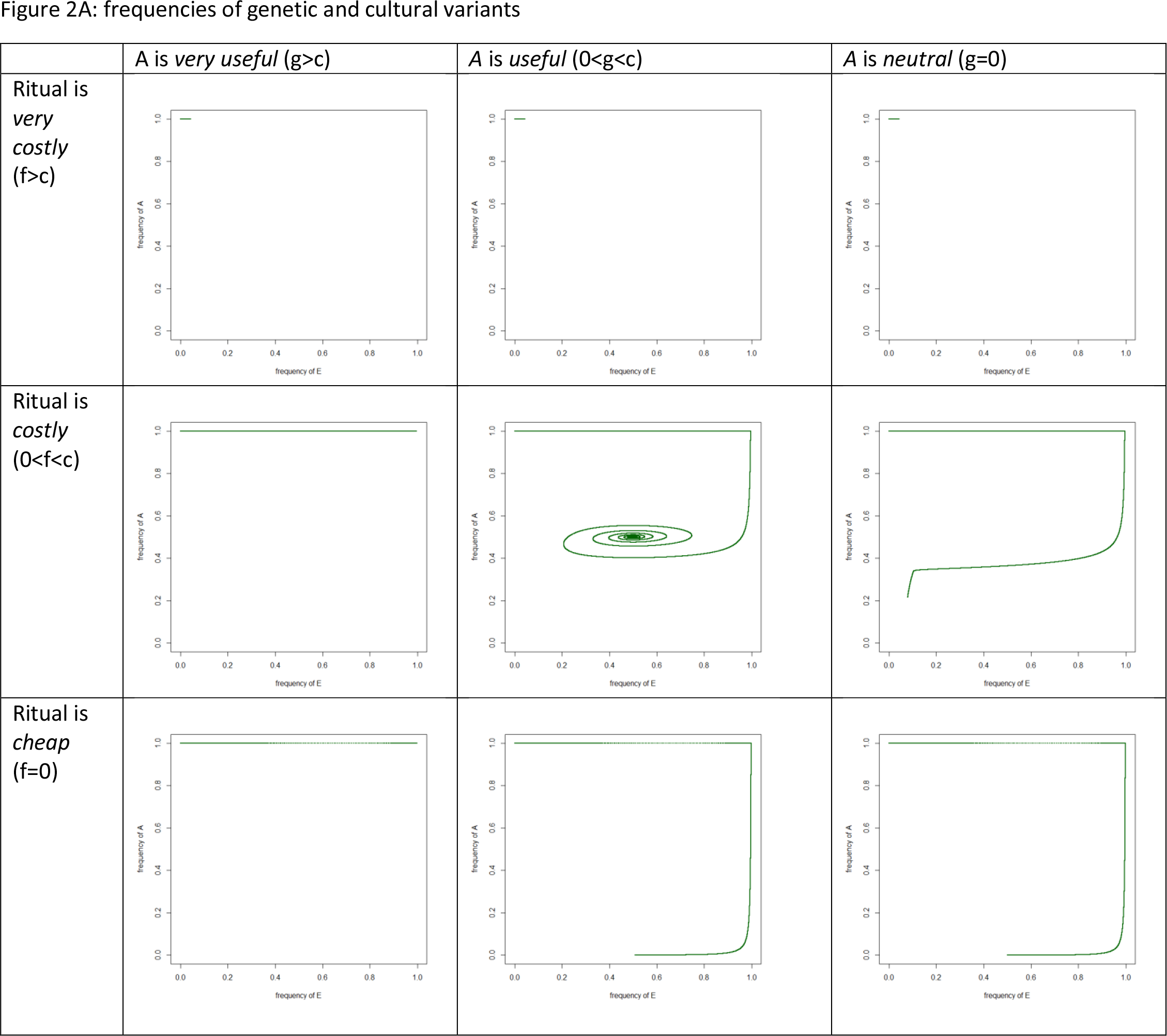
Simulation Results { b= 1, c=.5, w=10, Lifespan=50, mg = 2.5 × 10^−5^ sr= 1, sc = 0.3) Figure 2A: frequencies of genetic and cultural variants

**Figure 2B:**
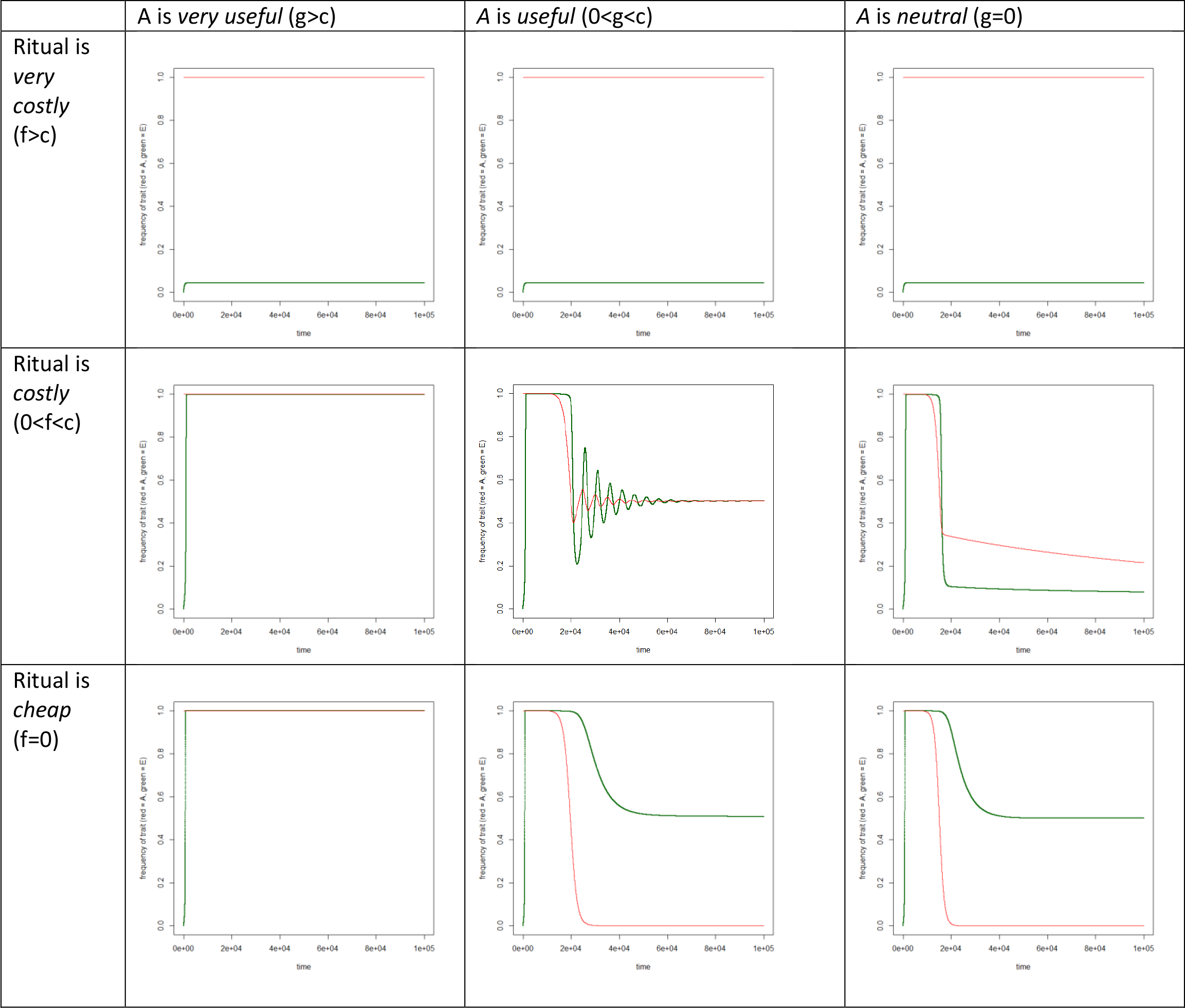
Gene frequencies over time

**Figure 2C:**
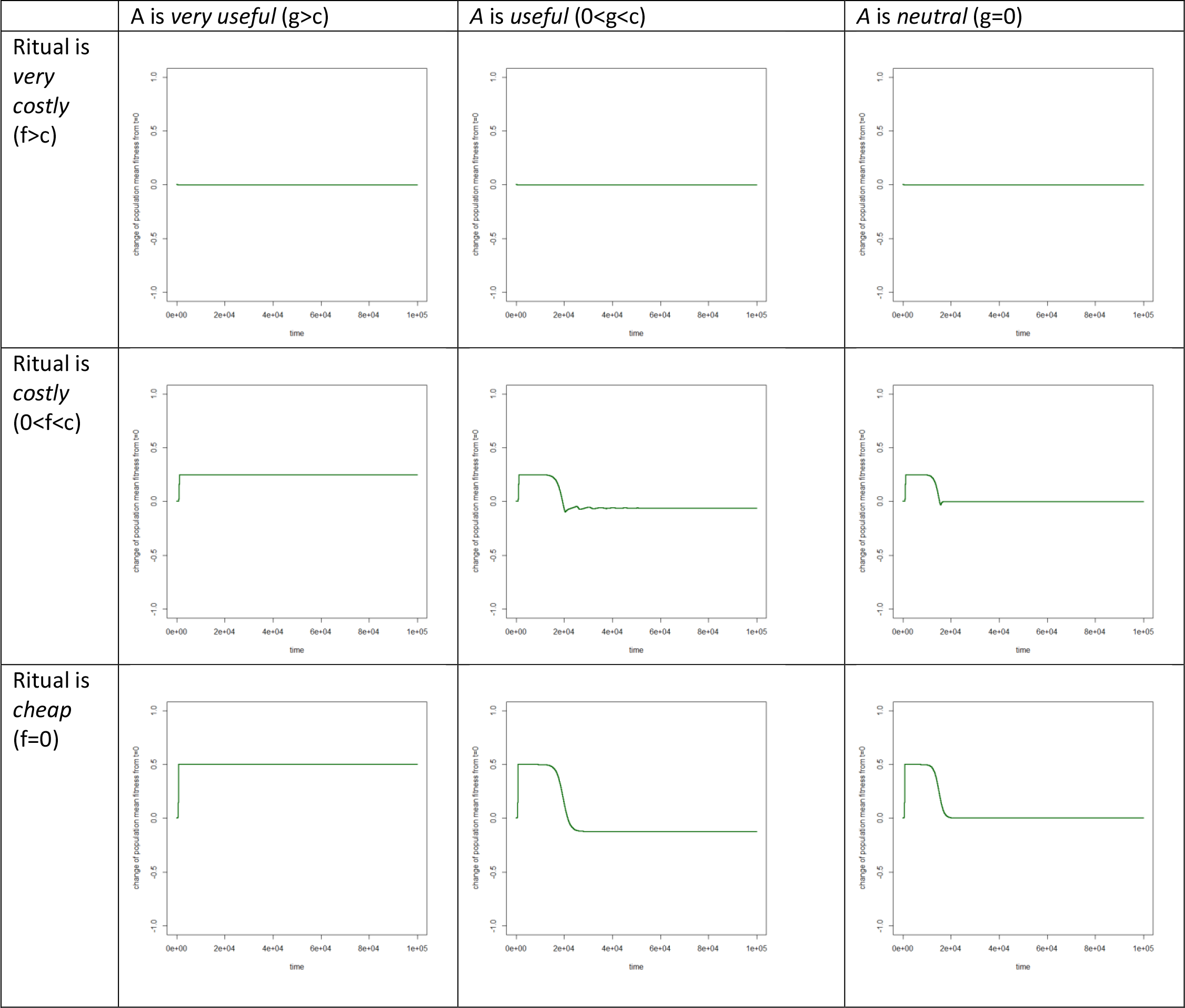
Population mean fitness over time

## Discussion and Conclusions

This analysis demonstrates how costly reciprocal rituals can, in some circumstances, hijack pre-existing, genetically based sensory biases to foster high levels of altruism in populations. Uncontroversially, if such sensory biases are pleiotropic with essential or very useful functions, then such rituals should become quite common and should regularly function to generate altruism. Also, when genes responsible for such responses are completely neutral otherwise and ritual is not very costly, then these genes will be replaced in the population with their neutral, non-prosocial alternative, a. This duplicates the results of green beard models. Different results occur for intermediate values of the benefit of the hijacked allele and cost of the ritual. These circumstances lead via damped oscillations to an equilibrium of mixed types. Even while eventual evolution of the population may lead to an elimination or reduction of the hijacked allele, the introduction of such a socially learned ritual may lead to long term stability of wide-spread altruism on the order of 10sof thousands of years … both a long time in terms of human recorded history and time enough within the history of human (and pre-human) social learning for gene-culture feedbacks to occur.

While this analysis replicates the results for equilibrium from a purely genetic model of coevolution, a number of differences can be seen comparing the simulations from the gene-culture coevolution model to simulations from a purely genetic coevolutionary model with otherwise identical structure (Frost, 2016a). These include

- As expected, for f<(b-c) (*costly* and *cheap* rituals, but not *too costly*) the frequency of E increases drastically faster at first than in the purely genetic case. In the case of the cultural model, this sweep to fixation happens in hundreds of years. In the genetic model, it takes about 35,000 years, despite being modeled on a much shorter lived species.
- Compared to the purely genetic model, the evolution of genetic trait *a* due to free-riding benefits happens much more rapidly. Because *E* moves to fixation so much faster in the gene-culture model, strong genetic pressure arises more quickly against allele *A*, causing the evolutionary change of this gene to be an order of magnitude faster than the response found in the gene-gene coevolution model. This is, again, despite the fact that the gene-gene coevolution model used a lifespan of 10 years instead of 50 years, which would otherwise result in evolutionary processes running 5 times faster, instead of an order or magnitude slower. Fast moving cultural evolution drastically accelerates genetic evolution. Similar rapidly rising, strong pressures on genes from fast-rising cultural traits has been observed with cattle raising and lactase persistence in Europeans (Itan, Powell, Beaumont, Burger, & Thomas, 2009).
- For the intermediate case of *useful* hijacked allele and *costly* ritual, the trait frequencies move through damped oscillations toward equilibrium. In both the genetic and gene-culture models, the damping of the oscillations and the range of the limit cycle is set by the rate at which traits randomly change, through mutation in the genetic case and innovation in the cultural case. In the purely genetic model, the population cycles widely amongst gene frequencies around equilibrium, unless there is a very high mutation rate. In the gene-culture model, the oscillations are rapidly damped until the population is quite close to equilibrium. IN the gene culture model, the damped oscillations of gene frequency is already within 1% of equilibrium within 50,000 years. In the genetic model, at 200,000 years, the populations is still in oscillations sweeping through most of gene frequency space. This difference is driven by the fact that the rate of random innovation is several orders of magnitude larger than the mutation rate. For exceptionally high mutation rates, the genetic model also cycles in toward equilibrium, albeit much more slowly, with more wide range cycles en route

The simulation results from the purely genetic model are included in an appendix.

For the case in which there is a stronger benefit for the hijacked trait than for cheating on the prisoner’s dilemma, the hijacking can be long term stable and lead to a higher population mean fitness. The non-prosocial gene can never invade and only exists as a rare allele maintained by mutation. Ritual performance sweeps to near fixation, and nonperformers of ritual persist only through personal innovation (in the cultural case) or mutation (in the case of genetically based rituals). This is independent of descent system and works as much for a genetic system as it does for a case of gene-culture coevolution.

Of course a long standing view of the problem with green beard effects, which this model resembles, is that there needs to be some sort of trait linkage for it to work. This does not seem obviously the case with this model, but one might ask if there is something analogous to trait linkage that is somehow slipped into assumptions of the model. The answer, I would argue, is ‘sort of’. There is something analogous to a trait linkage in the premise that A can be hijacked by a visible behavior (the ritual) to cause cooperation in the PD game. This assumption has the same kind of function as in the traditional green beard trait linkage assumption that the signal and altruism were coded by the same gene. In the case of this model, a single gene codes for prosocial reaction to ritual (as in synchronous movement practices) and to some other benefit (like parent child bonding via mimicry). This could be a pleiotropy, or it could simply be the same behavioral response to stimuli, but where the stimuli is generated through a novel socially learned behavior (the ritual). One could speculate about the possibility of an evolution of an ability to cheaply discern the two situations, to act like A in the original context, but like a in the PD game. Let’s call this allele A’. The reproductive value of A’E would then be

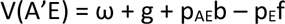

This at first seems like an obvious possibility. We can easily see the difference between our mother (with whom we should bond) and strangers (with whom we engage in the PD game). However, we are here modeling with allele A the genetically derived, emotion driven heuristics that constrain choices, not our culturally derived ability to distinguish these circumstances. Remember, before the advent of the ritual, there was no history of a need to make such a distinction on which genetic selection could act. Rather than being as simple as two different ‘person identification’ traits being separated by recombination, this would entail the genetic evolution of a new trait that suppressed the existing reaction to mimicry in the ritual. While possible and certainly evolutionarily advantageous in the absence of side effects, the genesis of such a new genetic trait via mutation is not by any means guaranteed or even expected to be likely. Of course, where this seems unlikely for the pure genetic model, it seems even less likely to be important for time scales of interest in the gene-culture interaction model, given the relative infrequency of positive random mutations.

It is interesting to note that while this model does allow for the evolution of levels of cooperation with unrelated others through ritual facilitation that would not otherwise be predicted, it does have the effect in some cases of lowering population-mean fitness despite the facilitated solution of cooperation problems. Specifically, where the high stakes benefits of cheating are greater than the benefits of the hijacked trait, and the ritual is costly, but less so than the net benefit of the PD, this will occur. Further, where the ritual is cheap, the hijacked gene, A, may eventually be almost entirely eliminated from the population. Ritual performance, E, becomes ubiquitous but has negligible effect on altruism.

These gene culture coevolution simulations give a number of different scenarios in which reciprocal rituals may be sustained in a population, even though costly. It may be that they take advantage of a regularly occurring sensory bias that is caused by relatively essential genes and thus regularly facilitate altruism. It may also be that they have done so in the past, but have caused a feedback process through gene-culture coevolution. In this case, there may be some genes associated with altruistic response to such rituals which have been under strong recent negative selection pressures, and it may be that altruism is less regularly facilitated by such rituals than they once were. It could be interesting, if one were able to find specific individual genes key to these prosocial responses, to test if there is evidence of recent selection on these genes.

As mentioned, this particular evolved psychological disposition, intention synchrony, is only one amongst many theories of ritual function. As an example of a quite different potential prosocial cultural hijacking, there is now a large body of empirical research which points to the prosocial benefits of calming ritual practices, like mindfulness meditation, yoga, or Tai Chi (Frost, in review) (Einolf, 2011) (Reb, Junjie, & Narayanan, 2010) (Hutcherson, Seppala, & Gross, 2008). Such calming rituals increase altruism, but do so generally, reducing parochialism. The cultural persistence of such rituals is not explained by this model, unless one adds other dynamics, such as positive assortment or secondary benefits of the ritual practice.

Humans are constantly innovating and experimenting with behavioral variation. Individual and social learning have the capacity to rapidly explore the design space of genetic constraints on behavior to find such idiosyncratic, genetically evolved prosocial responses and hijack them through artificial stimulation. Because of this rapid exploration of behaviors by individual innovation and experimentation, cultural hijacking of genetically evolved behavior for the solving of collective action problems is not only possible but arguably quite probable.

It is interesting to note that in the case of a mixed equilibrium, as the cost of ritual performance increases, the equilibrium frequency of both ritual performance and cooperation also increase. However, even while equilibrium population mean fitness increases with f, it is still less than the population mean fitness without the introduction of the ritual, unless g>c. In other simulations (not presented in this paper), in which different rituals were competed against each other, with all other things but ritual cost being equal, as long as f<b-c, the ritual with the larger cost (f) often eliminated the lower cost ritual from the population. I explored this question in more detail in the context of the genetic coevolution version of the model and found that the outcome of competing rituals very much depends on exactly how the different ritual forms interact with each other (Frost, 2016, appendix).

It seems odd that in the case of a mixed equilibrium, the equilibrium gene frequency has no dependence on the independent payoff, g, of the gene. This can be explained by the fact that the condition for fixation of both E and A is set by g. Specifically, as g>c, the population moves to relative fixation of both A and E, innovation and mutation notwithstanding. If g<c, then the equilibrium value of p_A_ is curiously independent of 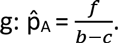. Similarly, the equilibrium value of P_E_ is independent of the cost of the ritual, 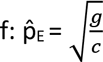

In the context of gene culture coevolution, socially learned rituals can arise and sweep to fixation in populations relatively quickly, leading to long periods of large genetic disequilibrium. This model then suggests one possible way that any genes vital for intention synchrony based bonding may have been under recent negative selection based on their being hijacked for synchrony rituals. Of course it may also be that, as synchrony-based rituals have a very deep history, this negative selection may have existed for long enough that gene frequencies have responded and this selection is now less.

The functional argument for the use of rituals to facilitate altruism often focuses on within-group altruism and is basically a cultural group selection argument (Rappaport, 1999; Wilson, 2002). Cultural group selection via evolved social norms may independently facilitate cooperation and altruism within groups through socially learned behaviors(Richerson et al., 2014). Where such dynamics occur, they may create independent feedback and positive selection on prosocial genes (Chudek & Henrich, 2011). This could then prevent or reduce the decline of genes responsible for such prosocial responses to ritual, and maintain high levels of ritual efficacy, even when the independent benefit of the gene is less than the potential advantage of free-riding (g<c in this model).

The purely genetic model indicates that this hijacking could happen by a genetically evolved ritual, and the phenomena may not be limited to gene culture interaction. This may be an explanation of social ritual actions observed in other animals like capuchin monkeys, chimpanzees, and birds (Perry et al., 2003)(Goodall, 2004)(Neuchterlein & Storer, 1982). Perhaps the inclination to perform ritual in these cases may be a genetic rather than socially learned development.

Finally, the model predicts that as f, the cost of performing the ritual gets small compared to (b-c), the coordination benefit in the PD game, the existence of the ritual virtually eliminates the prosocial disposition from the population. This is interesting because other cultural evolution models have argued for a positive effect of cultural evolution dynamics on prosocial genes (Chudek & Henrich, 2011). This suggests that gene-culture coevolution can effect genetically based prosociality in variety of counterintuitive ways. There is much modeling and empirical work to be done around this question.

## Appendix: R Code Gene Culture Coevolution Model

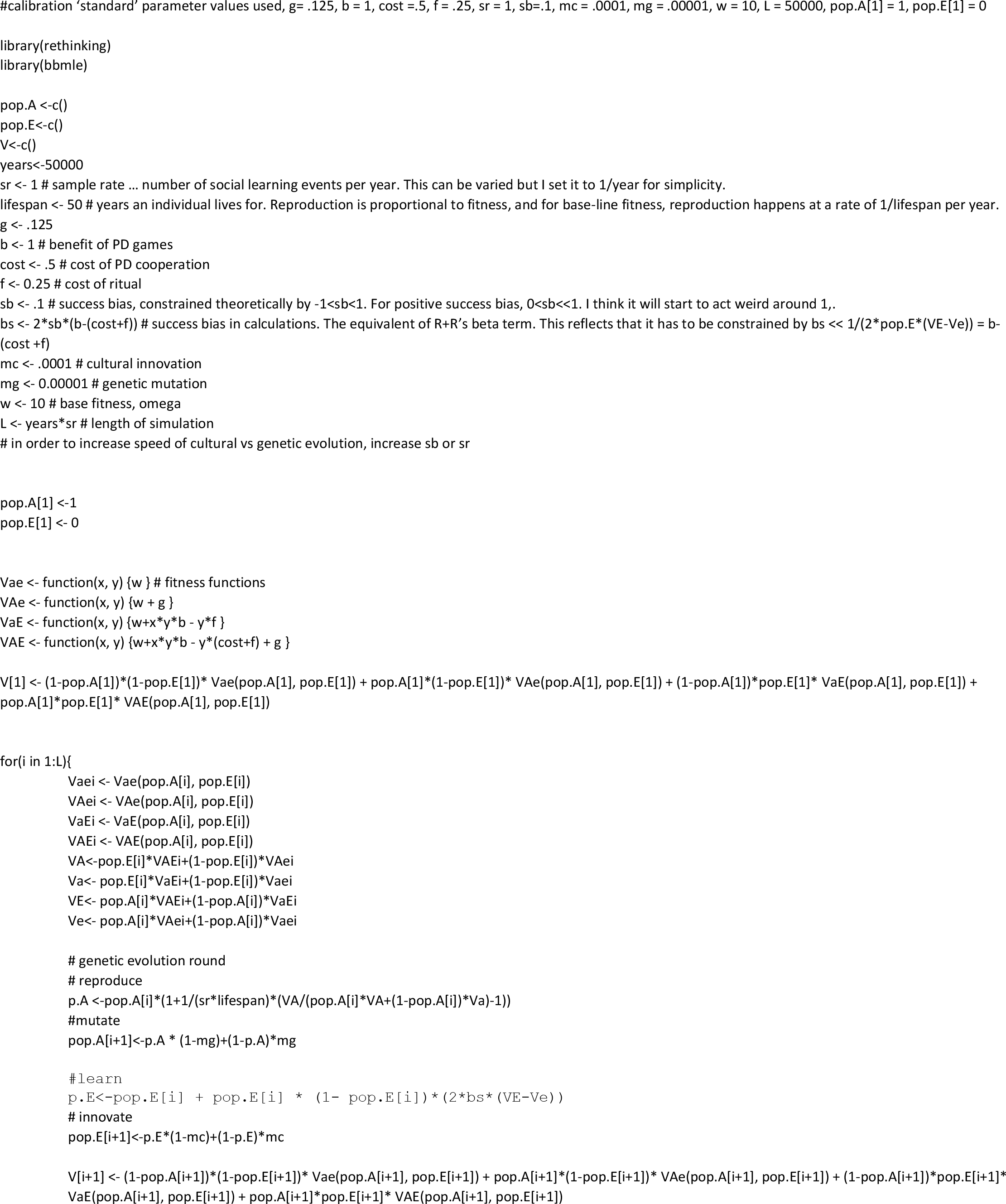
R Code Gene Culture Coevolution Model

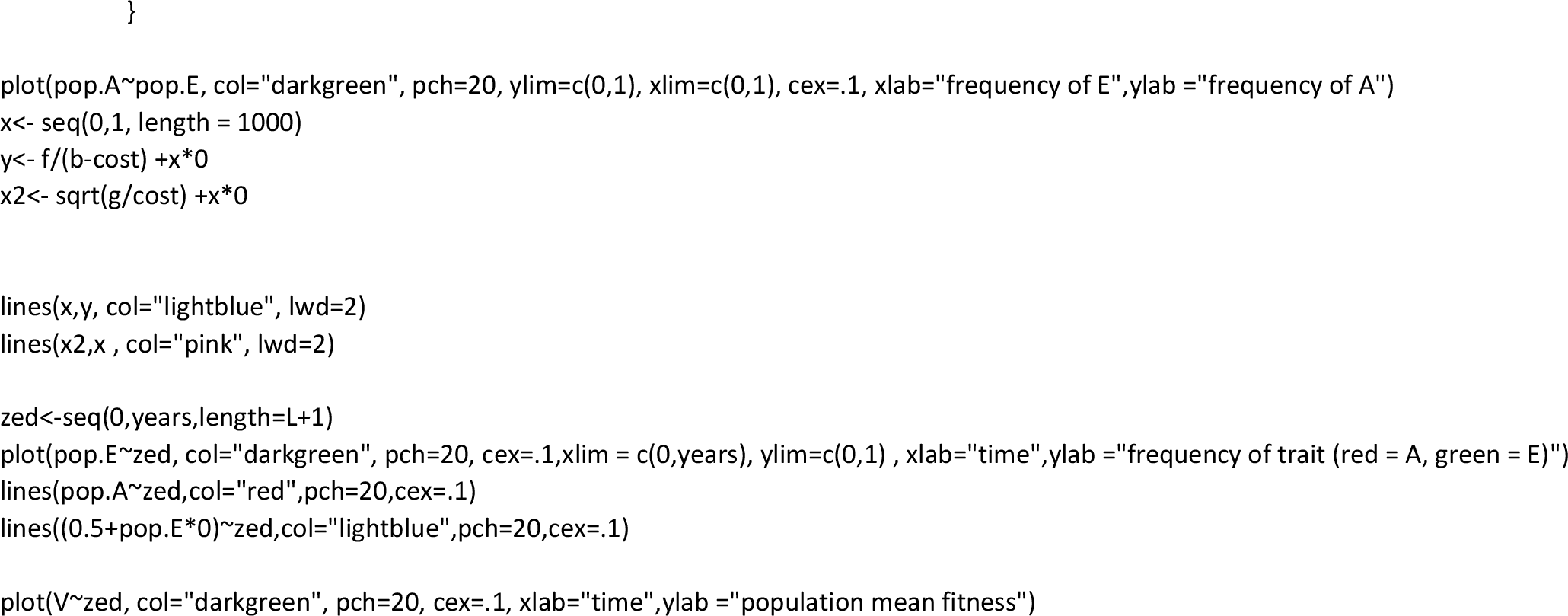
R code gene-gene coevolution, no stochasticity

Appendix: results from gene-gene coevolution simulations, copied from (Frost, 2016a)

**Figure 2:**
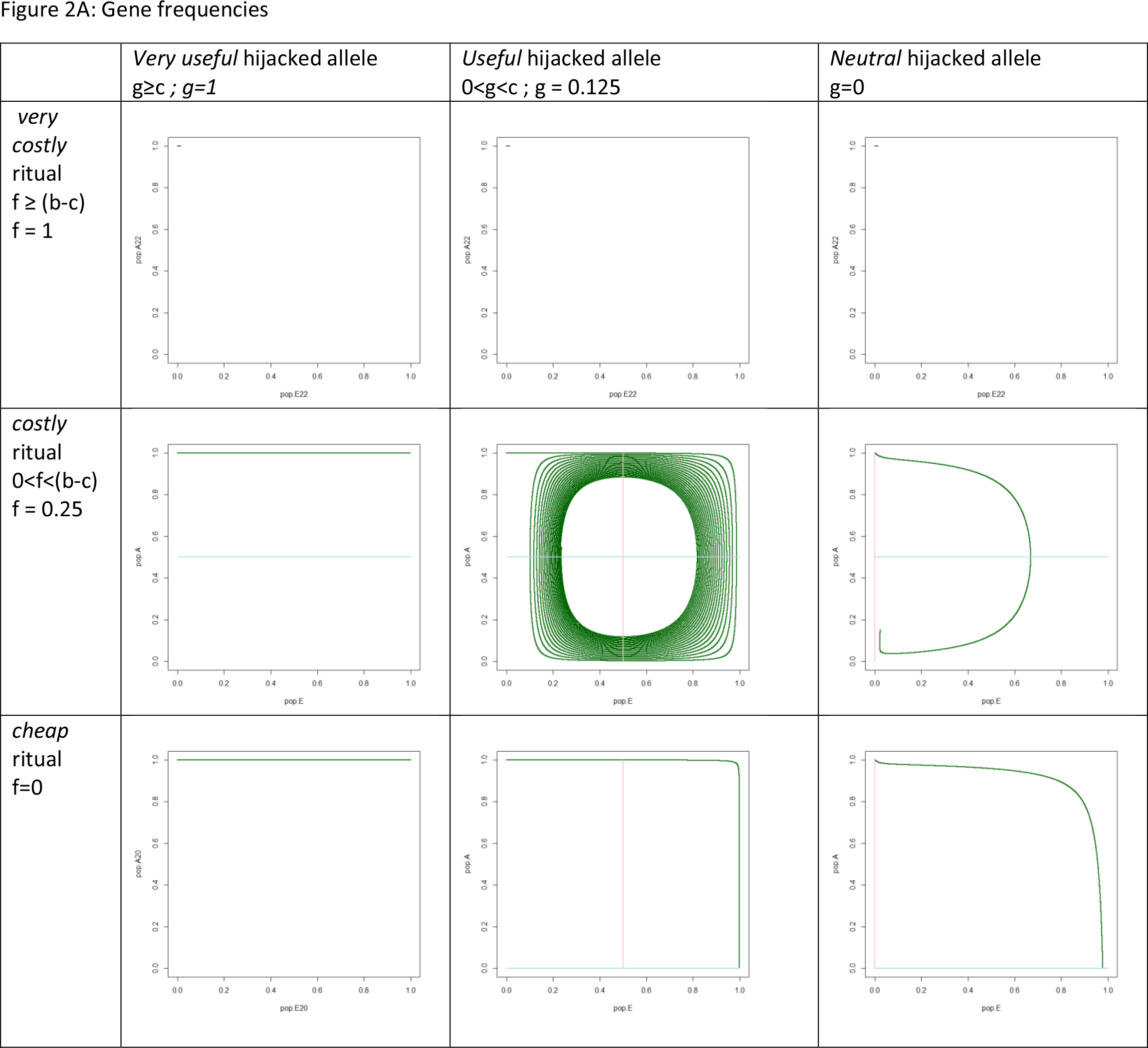
Simulation Results { b= 1, c=.5, w=10, r=.1, m = 1 x 10^−5^) Figure 2A: Gene frequencies

**Figure 2B:**
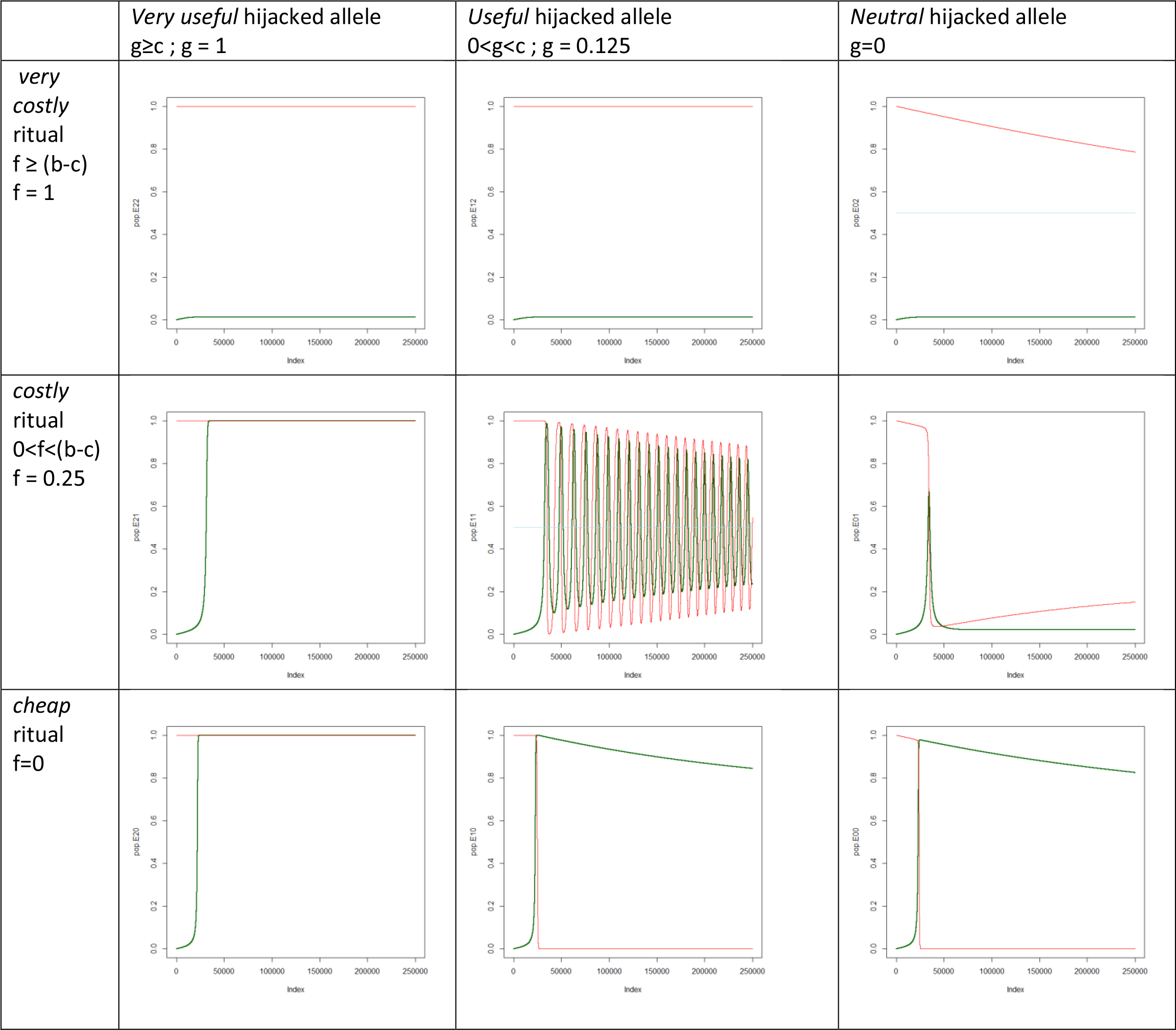
Gene frequencies over time

**Figure 2c:**
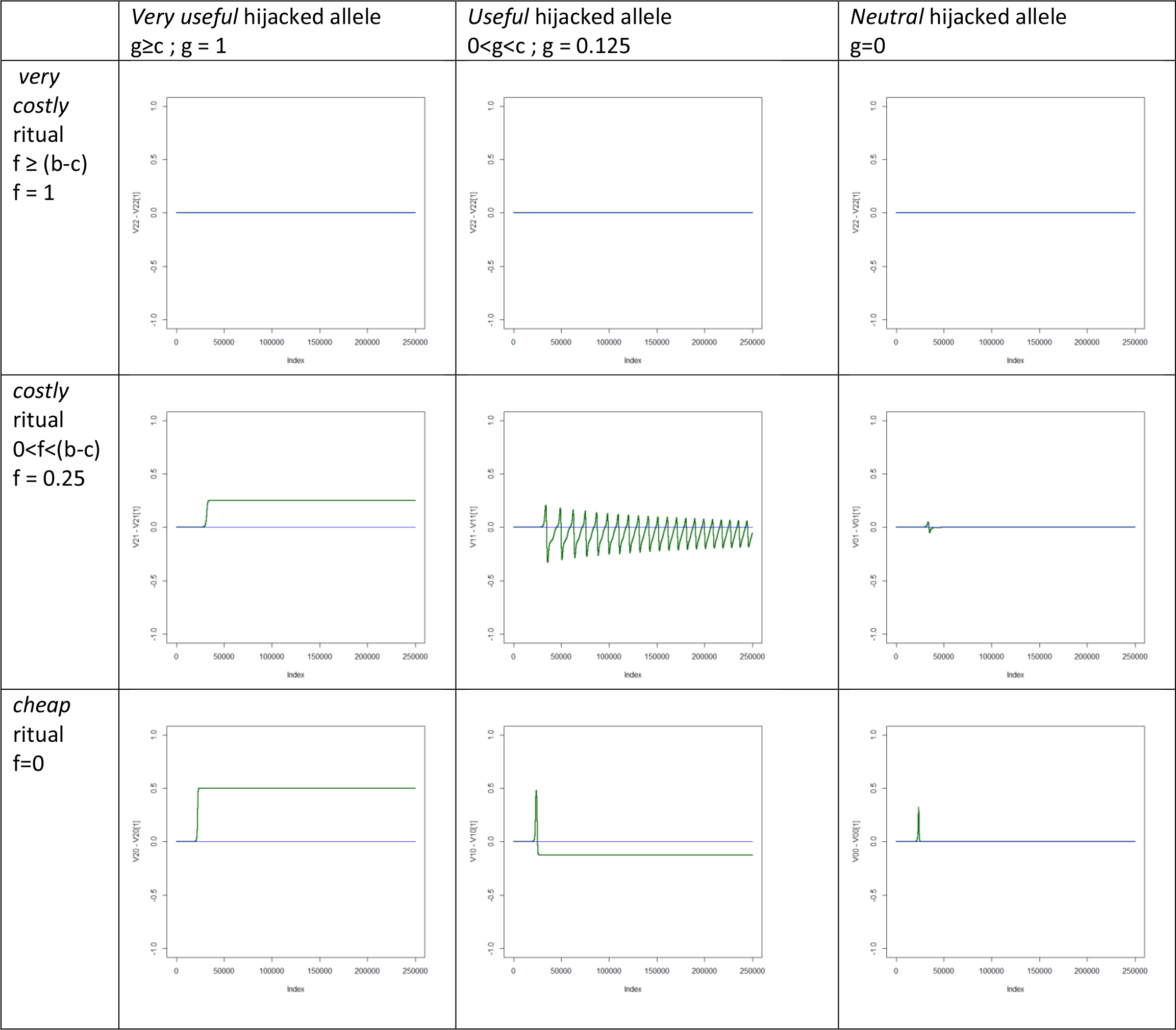
Population mean fitness over time

Appendix: Recombination-Gene Culture Coevolution model with Wright Fisher assumptions and covariance

The model used in this paper is a non-standard overlapping generations model. This was chosen to more realistically represent a human population. However, to make sure that the results of the simulations were not simply a fluke of the form of the model, the modelling exercise was repeated with a more commonly used form that assumed no overlapping generations.

Further, the assumption of complete recombination, while seemingly reasonable in the case of socially learned human behavior, is not completely accurate, and small amounts of gene-culture covariance have been shown to sometimes have large impacts on the trajectory of the population. While social learning, by its nature, leads to massive recombination between cultural variants and genes, it does not eliminate covariance. The model in this appendix models the recombination process explicitly and allows for covariance.

In this model, in each generation, the population reproduces in proportion to the fitness function, all adults die, and the population is normalized. In this process, cultural variants are past down vertically to offspring and the population undergoes both small, ecologically reasonable amounts of mutational change in the genes and small, reasonable rates of spontaneous innovation in culture, as with the model in the main text.

The population then goes through multiple rounds of horizontal, success-biased social learning in a well-mixed population, set here to 30 rounds. This would represent something like having a social learning opportunity once every year during adult life. The success biased social learning follows a typically used form, such that an individually encounters a random individual and decides to copy their socially learnable behavior with a probability equal to 0.5+βΔV, where β is a scalar for success bias (constrained to 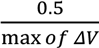) and ΔV is the difference of fitness between self and other.

The population then reproduces and the cycle repeats.

The simulation results of this model qualitatively reproduces the results of the model used in the body of the paper, as illustrated in Figure A1. “Index” in Figures A1B and A1C refers to number of generations for genetic reproduction.

Using reduced success biasing or using only one social learning round per generation slows the speed of the coevolutionary dynamics and moves the system in the direction of the gene-gene coevolutionary model, as illustrated in Figure A2. This makes sense, since the gene culture coevolution should have the same form as the gene gene coevolution, if social learning is only vertical social learning from parents.

**Figure A1:**
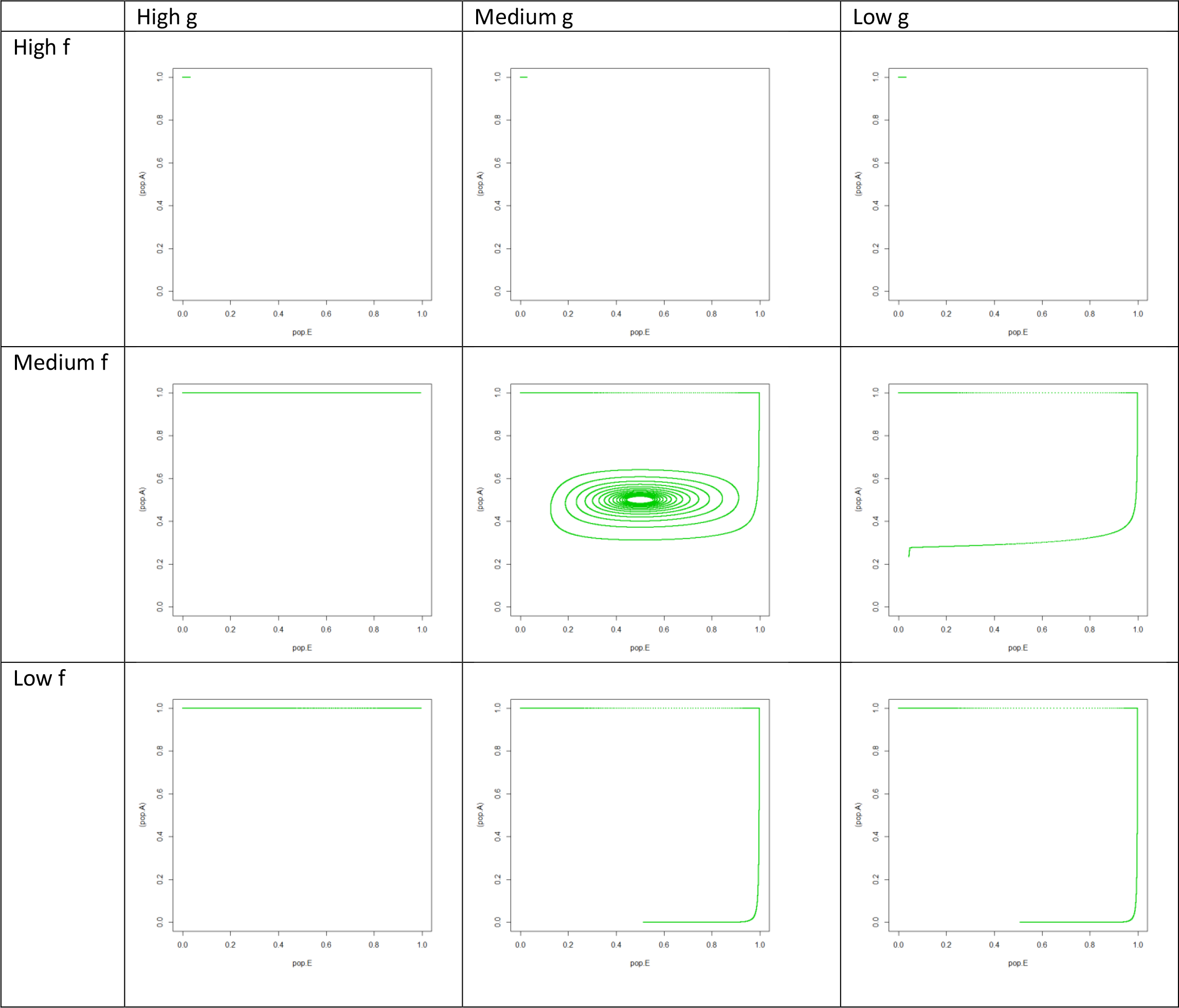
Simulation Results { b= 1, c=.5, w=10, r=.1, m = 1 x 10^−5^) Figure A1A: Gene Culture frequencies

**Figure A1B:**
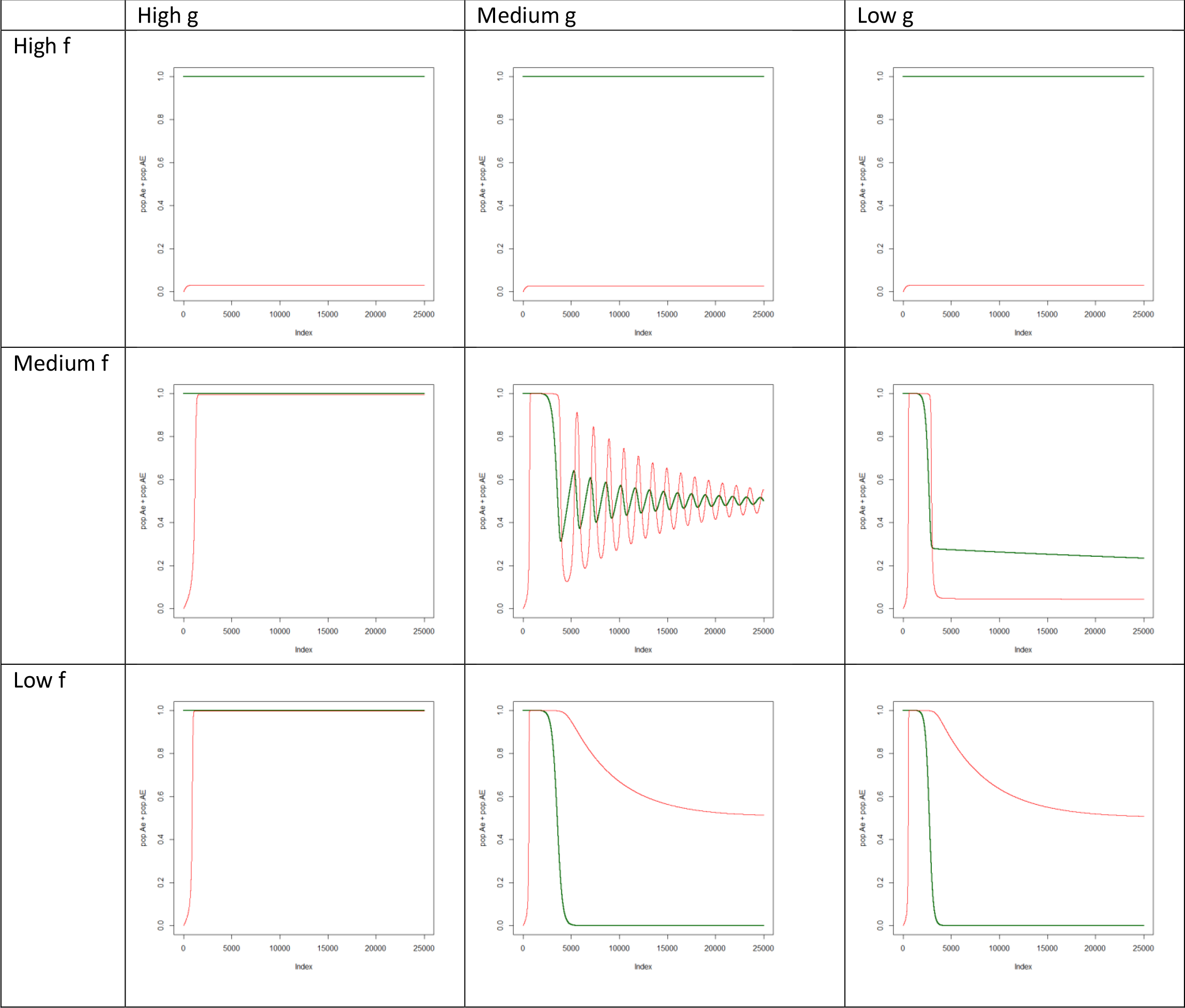
Gene and Culture frequencies over time

**Figure A1:**
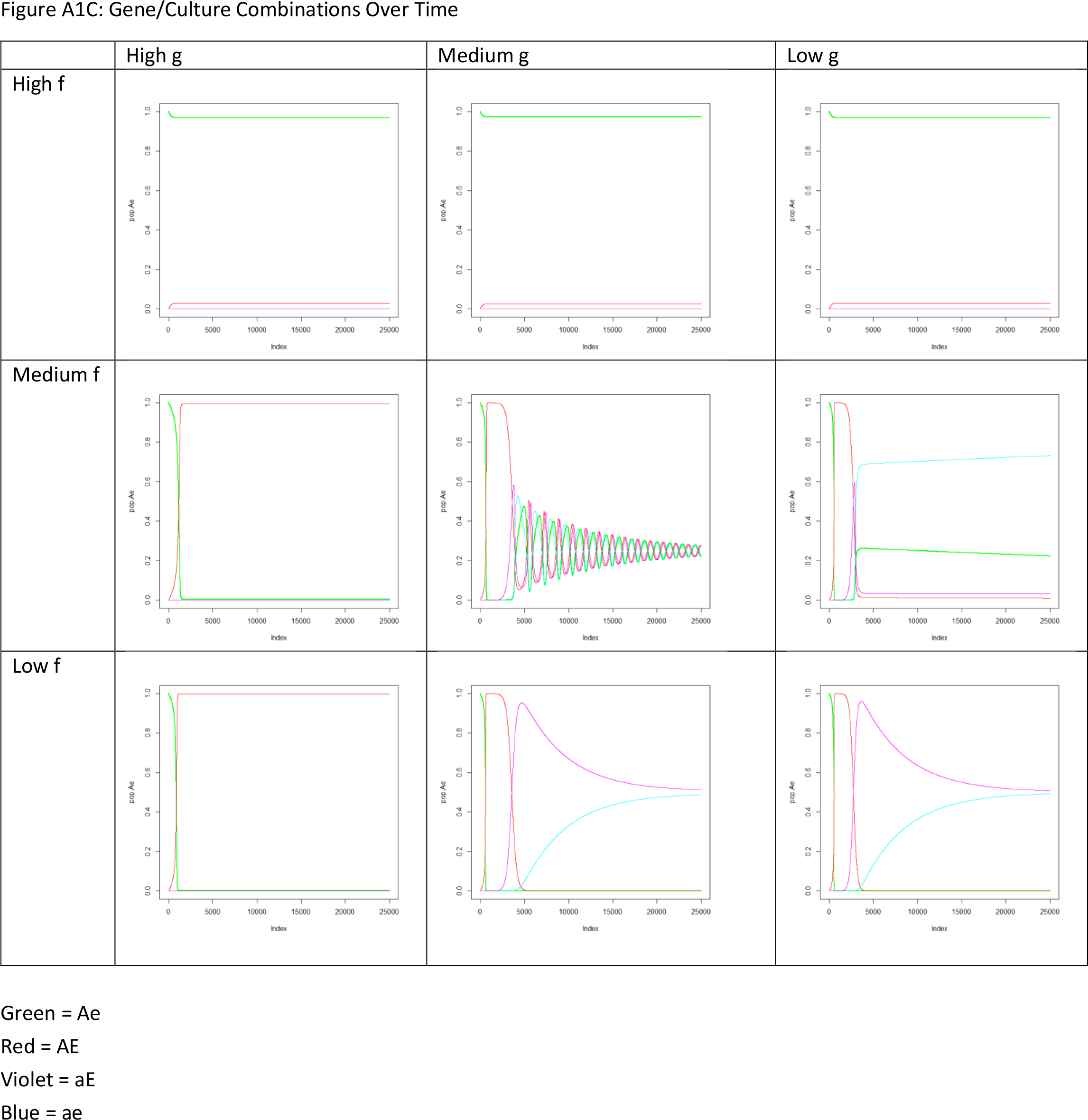
Gene Culture Simulation Results with no overlapping generations Figure A1C: Gene/Culture Combinations Over Time

**Figure A2:**
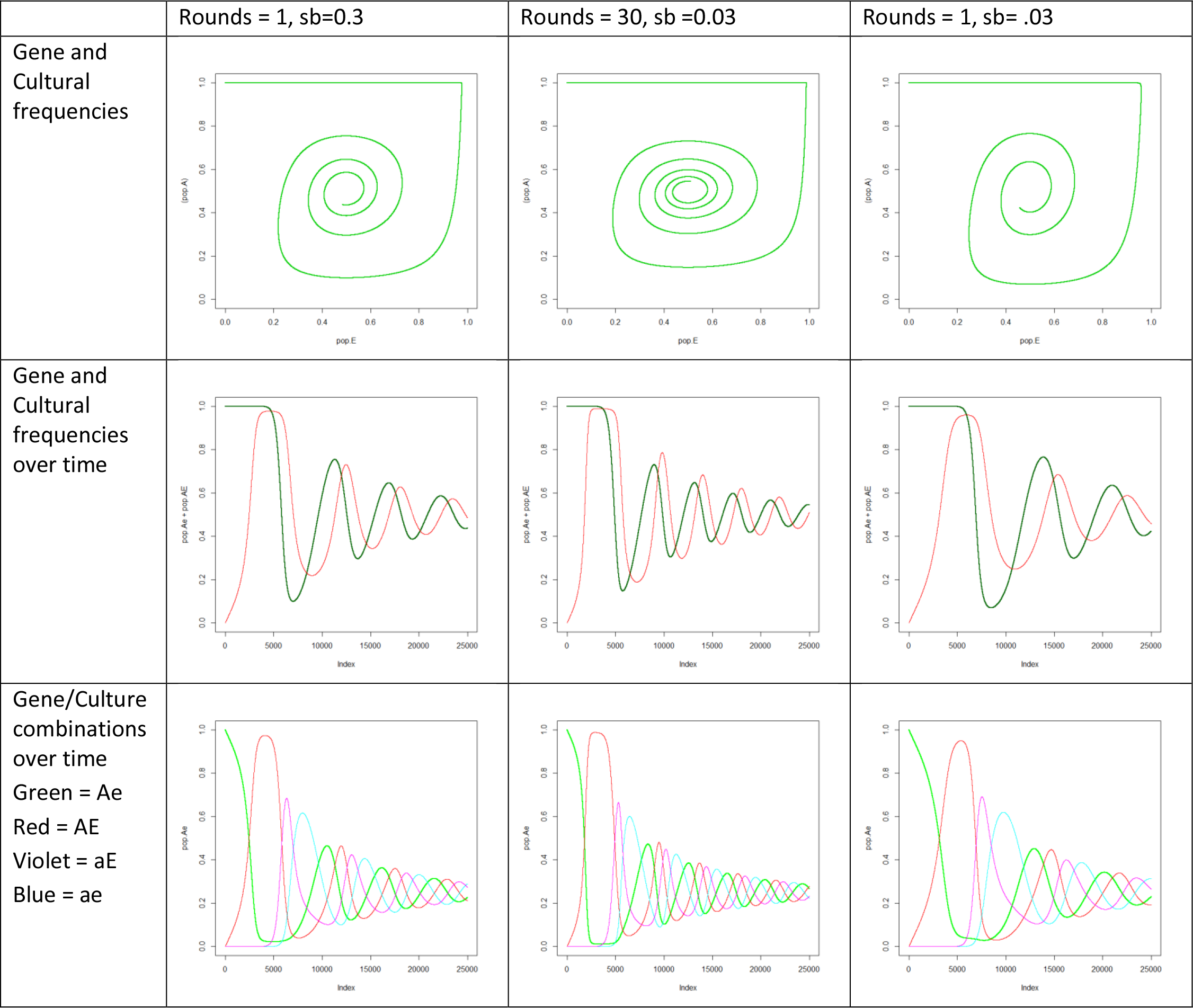
simulation of gene culture coevolution with smaller success bias and/or only one round of social learning per generation

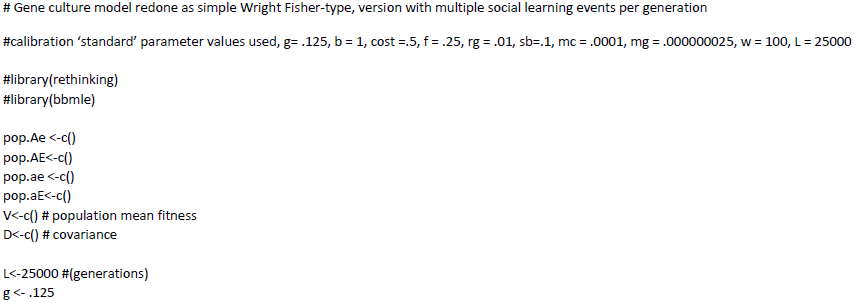

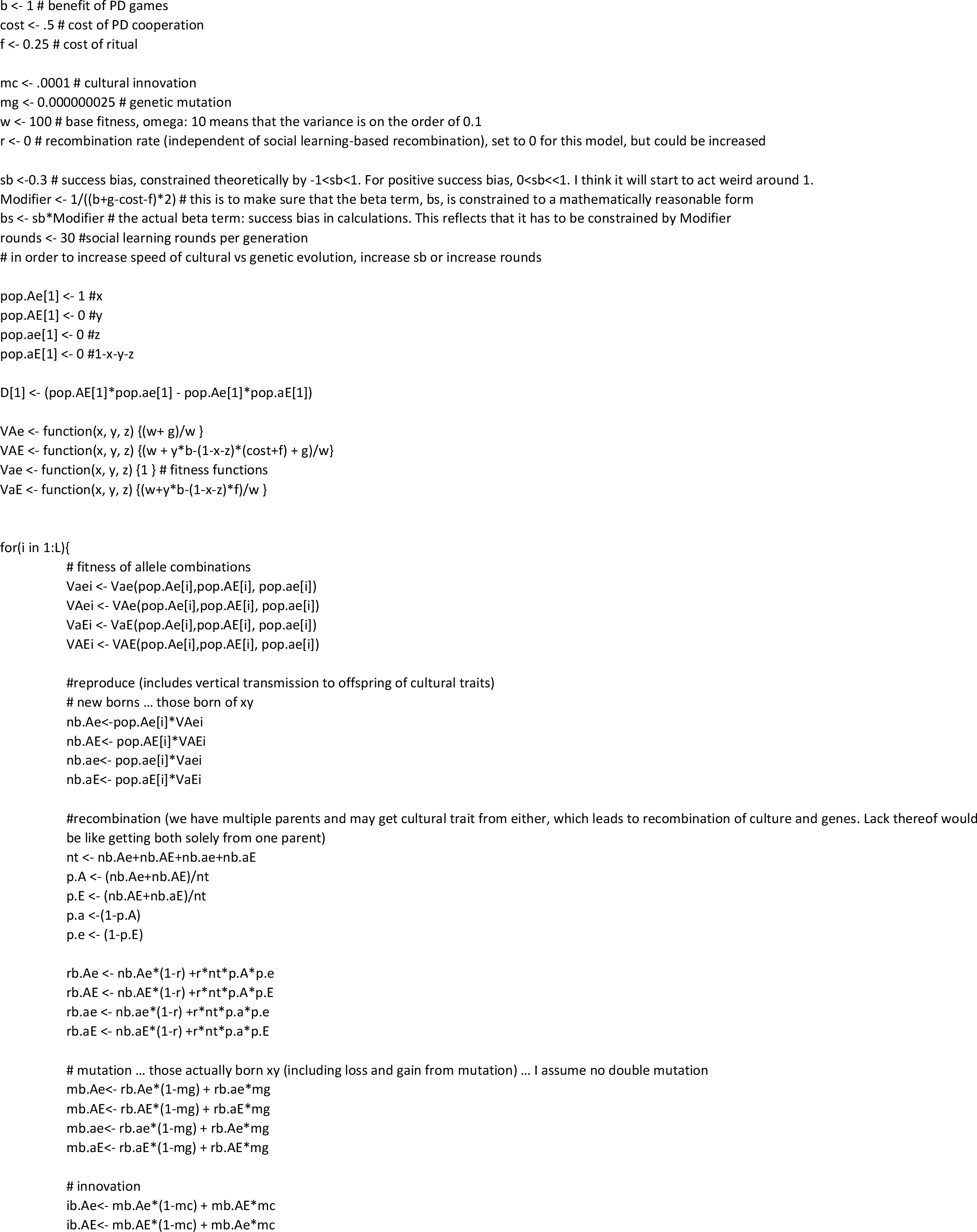

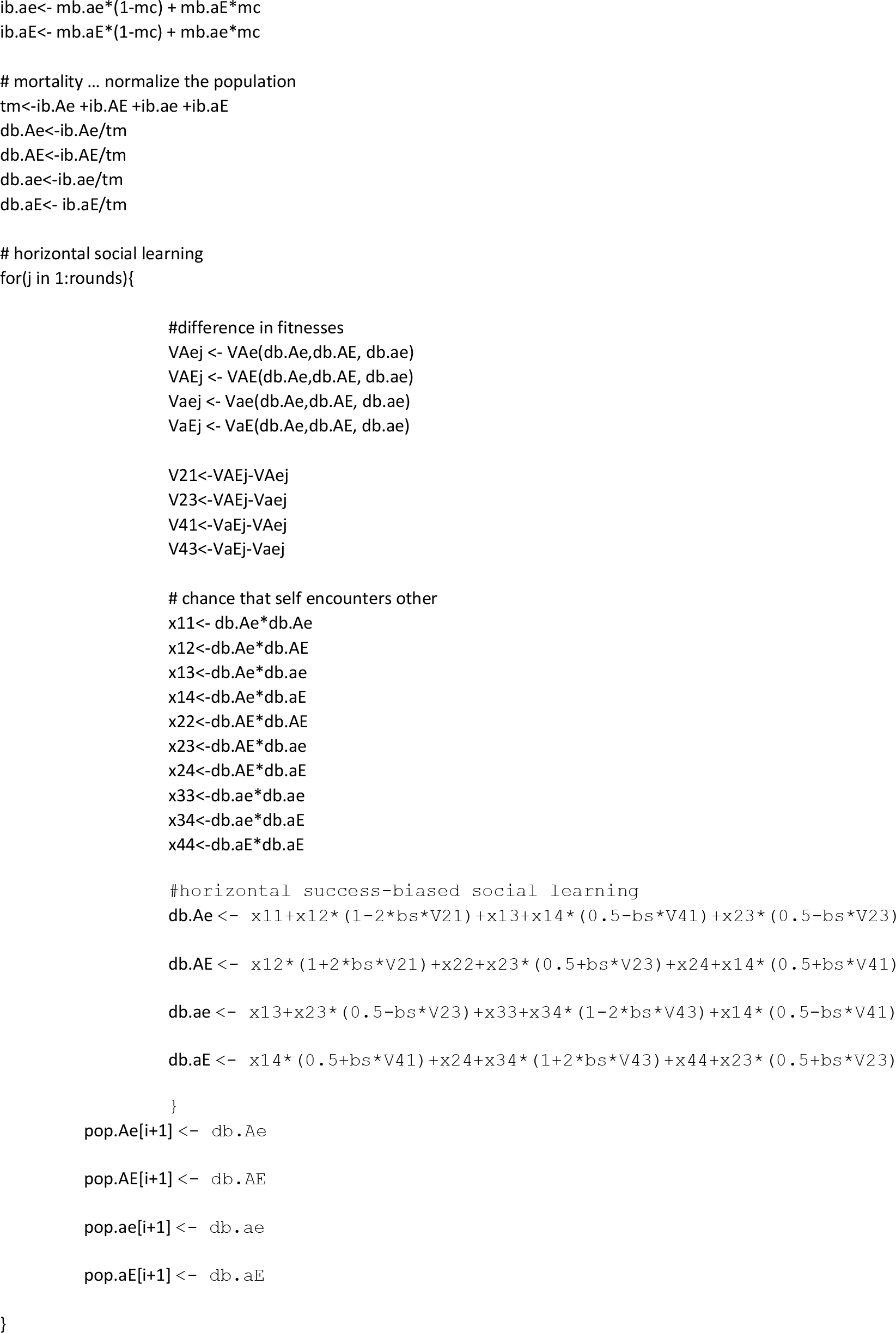

## Appendix 1: Innovation rate

Innovation, or individual learning, is the process of an individual deciding to adopt a behavior independent of social modeling of that behavior. An individual decides to experiment on their own with a novel attitude or behavior. In this case, an individual who does not practice ritual may invent the ritual behavior and offer to perform rituals with others, or it might mean that someone who does perform rituals may on their own decide to stop doing so. The latter seems quite plausible, as it is simply the loss of a socially learned behavior. The former at first seems less likely, but where socially learned ritual behaviors like dancing together are simply elaborations of instinctive ritual behaviors like unconscious mimicry of group partners, it becomes more plausible.

Individual learning is especially important as a single behavior moves towards fixation and models of alternative behaviors become rare. This allows for the regular re-introduction of rare behaviors in a population. When individual learning is a strong force, another potentially important effect of individual learning is to keep the frequency of specific behaviors from getting high. As is it is plausible that individual learning processes may be quite high at times, this may be an important consideration. Where there is gene-culture coevolution, this may in turn have effects on the rate of genetic evolution.

Figure A1 illustrates the effects of increasing the rate of individual learning on the trajectory of genetic and cultural variant frequencies over time. I use the scenario of *costly* ritual and useful hijacked allele. Other simulations in this paper use an individual learning rate of 1/1000 changes of behavior per year, symmetrically for both e to E and E to e. This assumption of symmetry is used for simplicity, but it is plausible that one direction or the other could be favored. For this appendix, two higher rates of invention are used: 1/100 and 1/10.

For i = 1/1000, ritual performance moves rapidly to fixation and gene frequency starts to significantly change after about 20k years. If the rate of invention is 1/100, the frequency of ritual performance never gets above 0.7, which results on a significantly decreased genetic fitness pressure against A. It eventually does decrease in frequency, but not until about 40k years and doesn’t get near equilibrium until about 80k years goes by. If instead there is a much stronger rate of individual invention, then the dominant force of random individual experimentation means that ritual frequency stays close to 50% and the selection pressure against A due to potential free-riding advantages never gets high enough. A stays at fixation and as a result cooperation induced by ritual behavior will be much higher that it would have been otherwise.

Of course in these simulation, a symmetric individual learning rate was used, unbiased by genetic fitness. It could very well be that there is a content bias for individual learning of ritual practice or non-practice that would establish a different frequency of ritual behavior. For 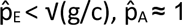. Thus, where content-biased individual learning is a sufficiently strong force relative to success biased social learning, keeping the frequency of *E* sufficiently low, *A* may remain at fixation and altruism may acheive higher frequencies than it might otherwise.

**Figure A1:**
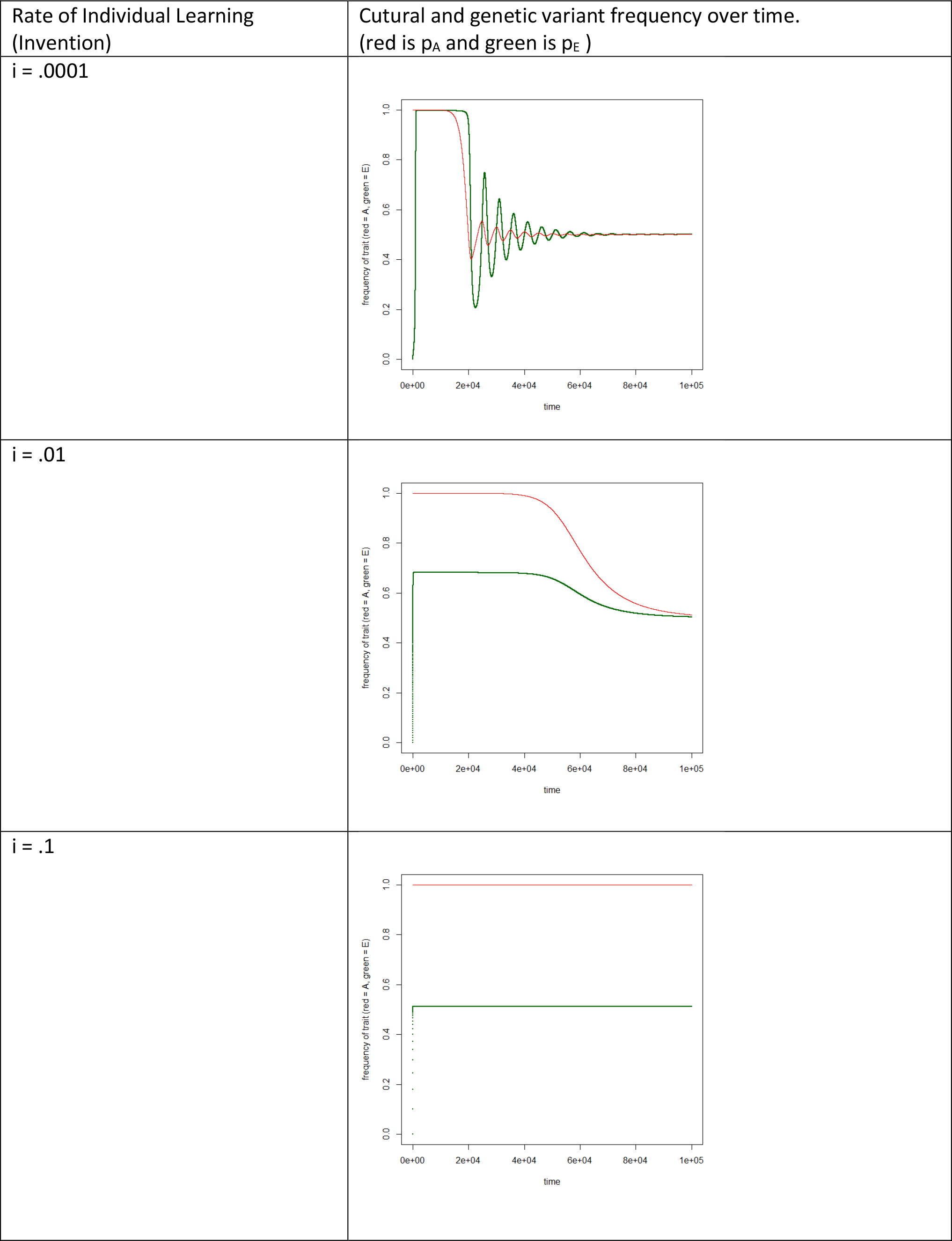
Gene Culture Simulation Results with no overlapping generations

